# Methylene blue, Mycophenolic acid, Posaconazole, and Niclosamide inhibit SARS-CoV-2Omicron variant BA.1 infection of human airway epithelial explant cultures

**DOI:** 10.1101/2022.03.30.486461

**Authors:** Romain Volle, Luca Murer, Anthony Petkidis, Vardan Andriasyan, Alessandro Savi, Cornelia Bircher, Nicole Meili, Lucy Fischer, Daniela Policarpo Sequeira, Daniela Katharina Mark, Alfonso Gomez-Gonzalez, Urs F. Greber

**Affiliations:** Department of Molecular Life Sciences, University of Zürich, Winterthurerstrasse 190, 8057, Zurich, Switzerland; Life Science Zürich Graduate School, ETH and University of Zurich, 8057 Zurich, Switzerland

**Keywords:** SARS-CoV-2 variant of concern Omicron (BA.1), Human nasal and bronchial airwayepithelialexplants, Drug repurposing, Methylene blue, Mycophenolic acid, Posaconazole, Niclosamide, Persistent infection

## Abstract

Sublineages of SARS-CoV-2 (severe acute respiratory syndrome coronavirus 2) Omicron variants continue to amass mutations in the spike (S) glycoprotein, which leads to immune evasion and rapid spread of the virus across the human population. Here we demonstrate the susceptibility of the Omicron variant BA.1 (B.1.1.529.1) to four repurposable drugs, Methylene blue (MB), Mycophenolic acid (MPA), Posaconazole (POS), and Niclosamide (Niclo) in post-exposure treatments of primary human airway cell cultures. MB, MPA, POS, and Niclo are known to block infection of human nasal and bronchial airway epithelial explant cultures (HAEEC) with the Wuhan strain, and four variants of concern (VoC), Alpha (B.1.1.7), Beta (B.1.351), Gamma (B.1.1.28), Delta (B.1.617.2) (**1, 2**). Our results not only show broad anti-coronavirus effects of MB, MPA, POS and Niclo, but also demonstrate that the Omicron variant BA.1 (B.1.1.529.1) sheds infectious virus from HAEEC over at least 15 days, and maintains both intracellular and extracellular viral genomic RNA without overt toxicity, suggesting viral persistence. The data underscore the broad effects of MB, MPA, POS, and Niclo against SARS-CoV-2 and the currently circulating VoC, and reinforce the concept of repurposing drugs in clinical trials against COVID-19.

## Introduction

The causative agent of COVID-19, SARS-CoV-2 rapidly evolves in the human population at high circulation frequency despite increasing natural and vaccine-induced immunity. Variants of concern (VoC) emerge and continue to turn over. For example, the Alpha and Beta VoC were first reported in the UK and South Africa in Summer 2020, and were replaced by the Delta VoC, first reported in India October 2020. Delta soon became the dominant VoC by mid 2021(**3**-**5**). Lately, the Omicron variant emerged, as first reported in South Africa in November 2021, and became abundant worldwide from December 2021 onward, replacing Delta by the beginning of 2022 (**6**). Omicron variants feature up to at least 30 amino acid substitutions, several deletions and also insertions in the spike protein (S) open reading frame (ORF). Remarkably, many of the substitutions are directly localized within the receptor-binding domain (RBD) (**7**), possibly the result of immune pressure upon infection and vaccination (**8**-**11**). Accordingly, neutralizing antibody titers against Omicron are found to be lower than against earlier VoC, as suggested in a preprint study from 23 laboratories (**12**). Nonetheless, Omicron appears to cause less severe disease than Delta, possibly because the immune status of many people still partially protects against Omicron and its VoC. However, Omicron continues to cause death notably affecting also non-vaccinated or incompletely vaccinated individuals (**11, 13**).

A growing body of evidence suggests that the multiple amino acid substitutions in the S-protein reduce the dependency of the virus on the serine protease TMPRSS2, compared to the Delta VoC (**14**). The altered usage of TMPRSS2 by Omicron appears to make the virion more dependent on low-pH endosomes and the cathepsin entry pathway (**14**-**16**). This highlights a considerable genetic flexibility of SARS-CoV-2 to adapt to different cell entry pathways, and may render the S-protein dependent entry a rather difficult target for powerful pan-interference strategies against SARS-CoV-2.

By engaging a multicycle drug repurposingscreen against coronaviruses we previously identified and validated several broadly acting compounds against SARS-CoV-2 infection of human nasal and bronchial airway epithelial explant cultures (HAEEC) grown at air liquid interface, namely, Methylene blue (MB), Mycophenolic acid (MPA), and Posaconazole (POS) (**2**).These compounds have been used in the clinics for unrelated applications, and can be considered for repurposing against SARS-CoV-2. Here we report that these compounds strongly inhibit SARS-CoV-2 Omicron, along with the anti-helminthic drug niclosamide (Niclo). Finally, we provide evidence for SARS-CoV-2 persistent infection of HAEEC.

### MB, MPA and POS reduce extracellular SARS-CoV-2 Omicron progeny levels

The partial immunity against circulating Omicron subvariants BA.1 and BA.2 and continuous evolution of SARS-CoV-2 necessitate the development of an arsenal of broad acting antiviral compounds. To address this need we tested the repurposing potential of several previously identified anti-SARS-CoV-2 compounds against the Omicron variant BA.1 (B.1.1.529.1), a close relative to the currently circulating BA.2VoC. The S-protein of BA.1 differs from the BA.2 S-protein by 12 amino acids, one nucleotide insertion and 5 nucleotide deletions (**17**).

We first focused on three compounds previously identified for their broad anti-coronavirus activity in cell culture, MB, MPA and POS (**2**). MB, MPA, and POS inhibit infection of primary human airway epithelial explant cultures (HAEEC) of nasal origin by the Wuhan strain, as well as Alpha, Beta, Gamma, and Delta VoC. Although their mode-of-inhibition is not known, these compounds do not inhibit SARS-CoV-2 cell entry, and have little effects on viral genome replication, but strongly inhibit the release of infectious progeny to the apical medium, indicating that they affect one or several post replication steps, such as particle formation or egress (**2**).

Nasal HAEEC (Epithelix, MucilAir™) grown at air-liquid interface (ALI) were apically inoculated with Omicron BA.1 (1,000 TCID_50_ per tissue) and treated with compounds added to the basolateral medium one day post infection (pi). Following five consecutive days of daily treatment and apical sampling, infectious virus titer was determined with 50% tissue culture infectious dose (TCID_50_) assays. Treatments with MB, MPA, and POS strongly inhibited the release of Omicron progeny (Fig. 1A), similarly as previously observed withthe Alpha, Beta, Gamma, and Delta VoC (**2**). The basolateral medium had no detectable virus titer, akin to the earlier study (**2**). The Omicron titers on the apical side, however, were in the range of 4.5 to 4.9-log_10_ TCID_50_/ml as early as one day pi (Suppl Fig. 1). MB, MPA, and POS reduced the average SARS-CoV-2 Omicron titers, albeit with different kinetics and efficiencies. Compared to the DMSO control-treated cells, MP, MPA or POS reduced the infection to 10 (±2.9), 36 (±18), and 58 (±31)%, respectively, at day one post treatment, and to 6.4 (±2.3), 18 (±4.3) 1.8 (±0.4)% at day five post treatment, notably without apparent tissue lesion, basolateral medium leakage or cell toxicity (Fig. 1B and 1C). Importantly, we could not detect evidence that MB and MPA induced phospholipidosis (Suppl. Fig. 2). Phospholipidosis manifests itself by a foamy appearance of internal cellular membranes likely owing to disregulated membrane signalling, sorting or transport (**18**). It was reported that cationic amphiphilic drugsmay have unspecific antiviral activity correlated with phospholipidosis, for example compounds such as chloroquine, that had been discussed for repurposing in the early days of COVID-19 (**19**).

**Fig 1.**
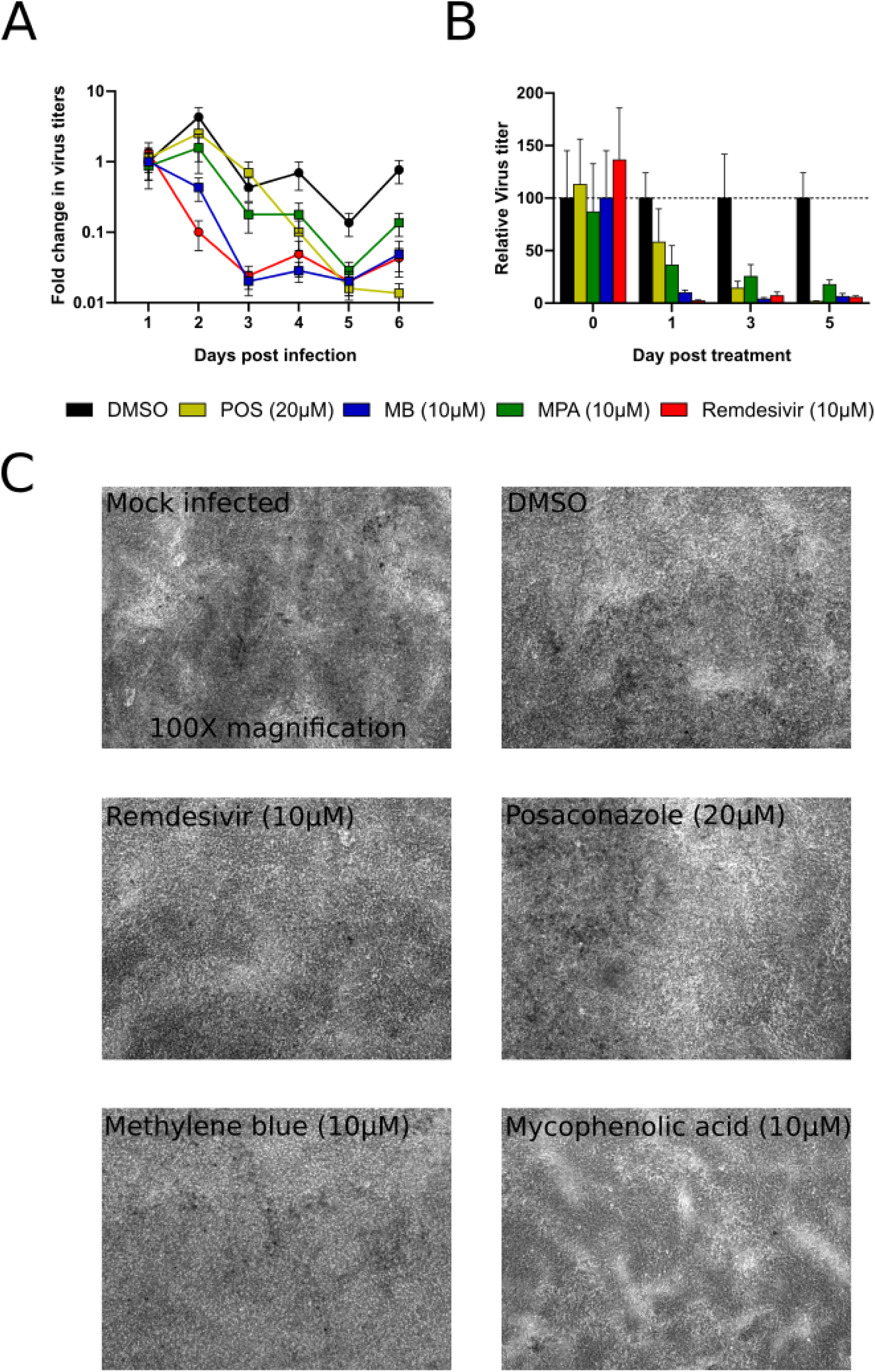
MPA, MB, and POS inhibit SARS-CoV-2 Omicron variant infection of nasal HAEEC. Antiviral effects of drug treatment represented as means ± SEM of three independent replicates. Nasal HAEEC grown at ALI were inoculated apically with 1,000 TCID_50_ units of SARS-CoV2 Omicron variant (day 0) and subjected to drug treatments in the basolateral medium, in a post-infection regimen starting at 1d pi. MB (10 μM), MPA (10 μM), and POS (20 μM) were administered daily until day 6. Remdesivir (10μM) and DMSO served as drug treatment controls. SARS-CoV-2 released at the apical side was collected daily by apical washing and quantified by TCID_50_ titration. A) Global fold change in virus titer. The baseline levels represent the apical means ± SEM of virus titer from the DMSO control sample at 1d pi (prior treatment). B) Relative change in virus titers of the treated inserts compared to the DMSO control C) Microscopic images of infected / treated nasal HAEEC at 6d pi. Pictures were taken through an inverted light microscope (Axiovert 135) at 100X magnification using CellF software (Olympus Soft Imaging Solutions GmbH, version 3.0).

### Comparable growth of SARS-CoV-2 Omicron in nasal and bronchial HAEEC

We next compared the susceptibility of human primary epithelial explant cells of nasal and bronchial origin to Omicron BA.1 infection. Our results indicate that Omicron similarly infected both nasal and bronchial lung epithelial cells yielding apical titers in the range of 10^4^ to 10^5^ TCID_50_/ml (Fig.2). These results suggest that bronchial cells can be used as a model for drug assessment against SARS-CoV-2 Omicron, and are in agreement with a report using *ex vivo* cultures, where Omicron exhibited higher replication in bronchial cells than in lung parenchymal cells mimicking alveoli of the lower respiratory tract (**16**).

**Fig 2.**
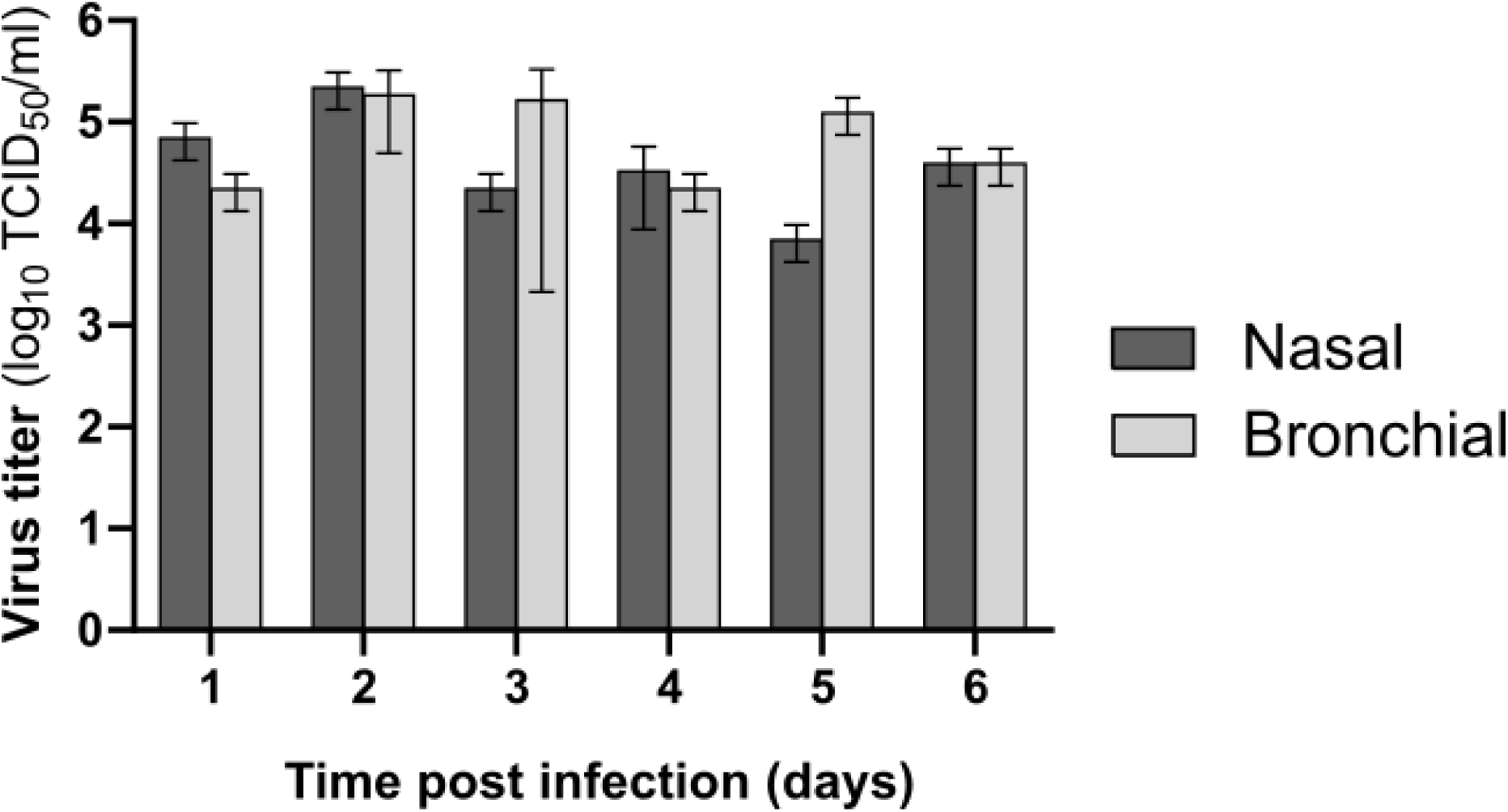
SARS-CoV-2 Omicron infection of nasal and bronchial HAEEC. Nasal and bronchial HAEEC grown at ALI were inoculated apically in duplicate with 1,000 TCID50 units of SARS-CoV2 Omicron. Virus released at the apical side was collected daily by apical washing and quantified by TCID_50_ titration. Data are represented as means ± SD of virus titers.

### Niclosamide inhibits SARS-CoV-2 Omicron infection of bronchial HAEEC

To further address theneed of acute medical treatment options with broadly available, safe and effective antivirals, we tested if Niclo inhibited Omicron infection of bronchial HAEEC. Niclo is FDA-approved for the treatment of tapeworm infections. It acts as a protonophore and has broad anti-helminthic and antiviral activity owing to its ability to neutralize acidic cytoplasmic membrane compartments, as well as its effects on membrane trafficking and cell signaling processes (**20**-**23**). Recently, Niclo was shown to inhibit infection of primary human bronchial epithelial cells with SARS-CoV-2 Alpha, Beta, and Delta VoC (**1**).

To test if Niclo affected SARS-CoV-2 Omicron infection, we apically inoculated Omicron onto bronchial HAEEC, and treated the cells with different concentrations of Niclo (20, 10, 5, and 1µM) one day pi for up to day 8 in daily treatments and apical sampling, followed by TCID_50_ titration of infectious progeny production. While 1µM of Niclo had no effect on Omicron, and 20µM was toxic for the cells, a daily treatment with intermediate concentrations of 5 or 10µM reduced the infectious titer of Omicron at days 3 to 8 up to 2 log_10_ (Fig.3). Similarly we could not detect evidence that Niclo induced phospholipidosis (Suppl. Fig. 2). These results were similar to the ones with the Alpha, Beta, and Delta VoC reported earlier (**1**). Notably, our effective concentrations of Niclo were slightly higher than those used by Weiss et al., namely 5-10µM versus 1.25-5µM. This difference possibly reflects the post-exposure treatment in our case, versus the preexposure treatment by Weiss and colleagues, or alternatively, donor-to-donor variability of the HAEEC. Cell-to-cell and donor-to-donor variability can have a significant influence on the infection outcome both *in vivo* and *in vitro* (**24**-**26**). Taken together, Niclo may be considered as a potential inhibitor of SARS-CoV-2 with broad effects on VoC. Not surprisingly though considering the low systemic availability of Niclo (**27**), a recent phase 2 randomized clinical trial with per oral application of Niclo did not significantly reduce the period of contagious SARS-CoV-2 infection in a cohort of 33 patients compared to a placebo cohort of 34 patients (**28**). However, aerosolized formulations of Niclo may be worth testing against COVID-19, as they can be safely applied to human airways (**28, 29**).

**Fig 3.**
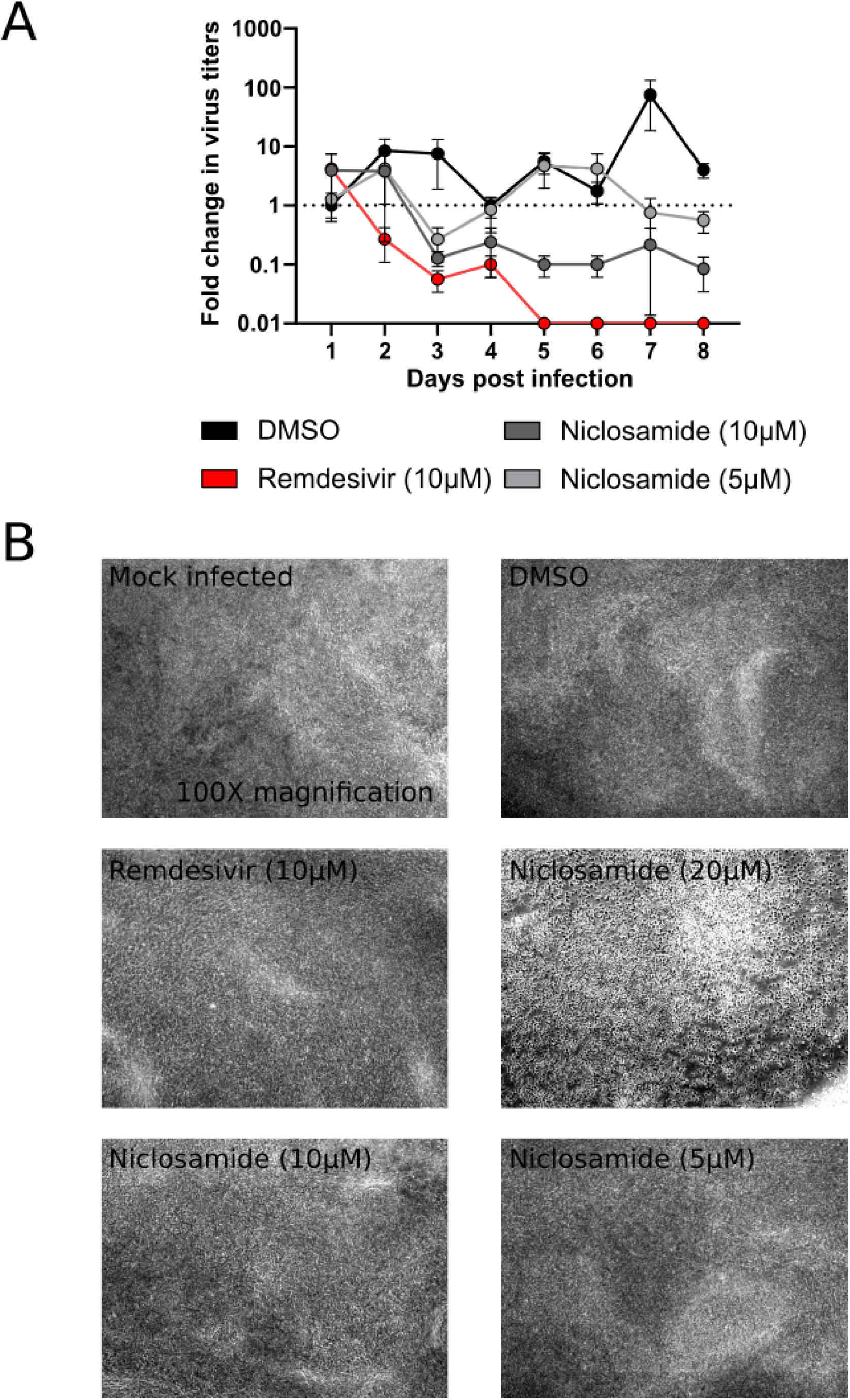
Niclosamide inhibits SARS-CoV-2 Omicron infection of bronchial HAEEC. Bronchial HAEEC grown at ALI were inoculated apically with 1,000 TCID_50_ units of SARS-CoV2 Omicron (day 0), and treated daily with Niclo in the basolateral medium starting at 1 d pi. Remdesivir (10 μM) and DMSO served as positive and negative control treatments. A) The antiviral effects of Nicloare represented as means ± SEM fold change in virus titers of two independent replicates. Baseline levels represent the apical means ± SEM of virus titer inthe DMSO control sample at 1dpi. B) Microscopic images of infected / treated nasal HAEEC at 6d pi. Pictures were taken through an inverted light microscope (Axiovert 135) at 100X magnification using CellF software (Olympus Soft Imaging Solutions GmbH, version 3.0).

### Persistent infection of nasal HAEEC by SARS-CoV-2

Although the origin of Omicron variant is still debated, a chronic infection has been a distinct possibility. An early report of the COVID-19 Genomics Consortium UK (**30**) hypothesized that chronic infections may have played a role in the origin of the Alpha variant (B.1.1.7). This may not be far-fetched because human coronaviruses have been long known to establish and maintain persistent infections *in vitro*. For example, the alpha coronavirus CoV-229E maintains a persistent infection of human fetal lung cells (L132) for up to 300 passages over a period of two years and produces infectious progeny (**31**). The beta CoV-OC43 persists in infected neurons, astrocytes, microglial, and oligodendrocytes cell lines andwas shown to release infectious particles for up to 25 passages andmore than one hundred days (**32**).

The question if SARS-CoV-2 persists *in vivo* has been debated, partly, because it is difficult to discriminate between a persistent infection and a second infection. A case report from South Africa, however, provided evidence that a 22 years old, HIV-positive woman under anti-retroviral therapy was persistently infected with SARS-CoV-2 (**33**). Over the course of 9 months, the virus acquired 21 nucleotide mutations, including 10 in the spike coding sequence leading to the substitution of six amino acids in the receptor binding domain (RBD), the deletion of three amino acids in the N-terminal domain, and the substitution of two amino acids in the S2 subunit. In addition, a 45 years old immunocompromised man produced infectious SARS-CoV-2 for up to 150 days, and was monitored by whole-virus genome sequencing which revealed evidence for fast continuous viral evolution (**34**). Yet another example for long term intra-host evolution and SARS-CoV-2 persistence was recently reported with a diabetic male patient with Non-Hodgkin Lymphoma (**35**). Additionally, a study with 203 post-symptomatic patients showed evidence for SARS-CoV-2 persistence, as indicated by RT-qPCR in pharyngeal samples from 26 individuals at 15-44 days and from 5-individuals at 85-105 days post recovery (**36**).

We took apical samples from the Omicron infected, DMSO control nasal HAEEC cells (presented in Fig. 1) for up to 15 days, and found a continuous titer between about 10^4^ and 10^5^ TCID50/ml in the apical milieu (Fig. 4A). The RT-qPCR genome equivalents were between about 10^7^ and 10^9^ copies / ml, indicating continued production and release of viral components over several weeks. In accordance, infected nasal HAEEC cells fixed at 7d and 21d pi followed by RNA FISH staining demonstrated the presence of intracellular SARS-CoV-2 ORF1ab RNA(+) fluorescent puncta predominantly in the cell layer near the apical side of the pseudo-tissue (Fig. 4B). Similarly, the 21d infected HAEEC cells also exhibited a clear staining of the SARS-CoV-2 ORF1ab RNA(+), albeit to a lesser extent.

**Fig 4.**
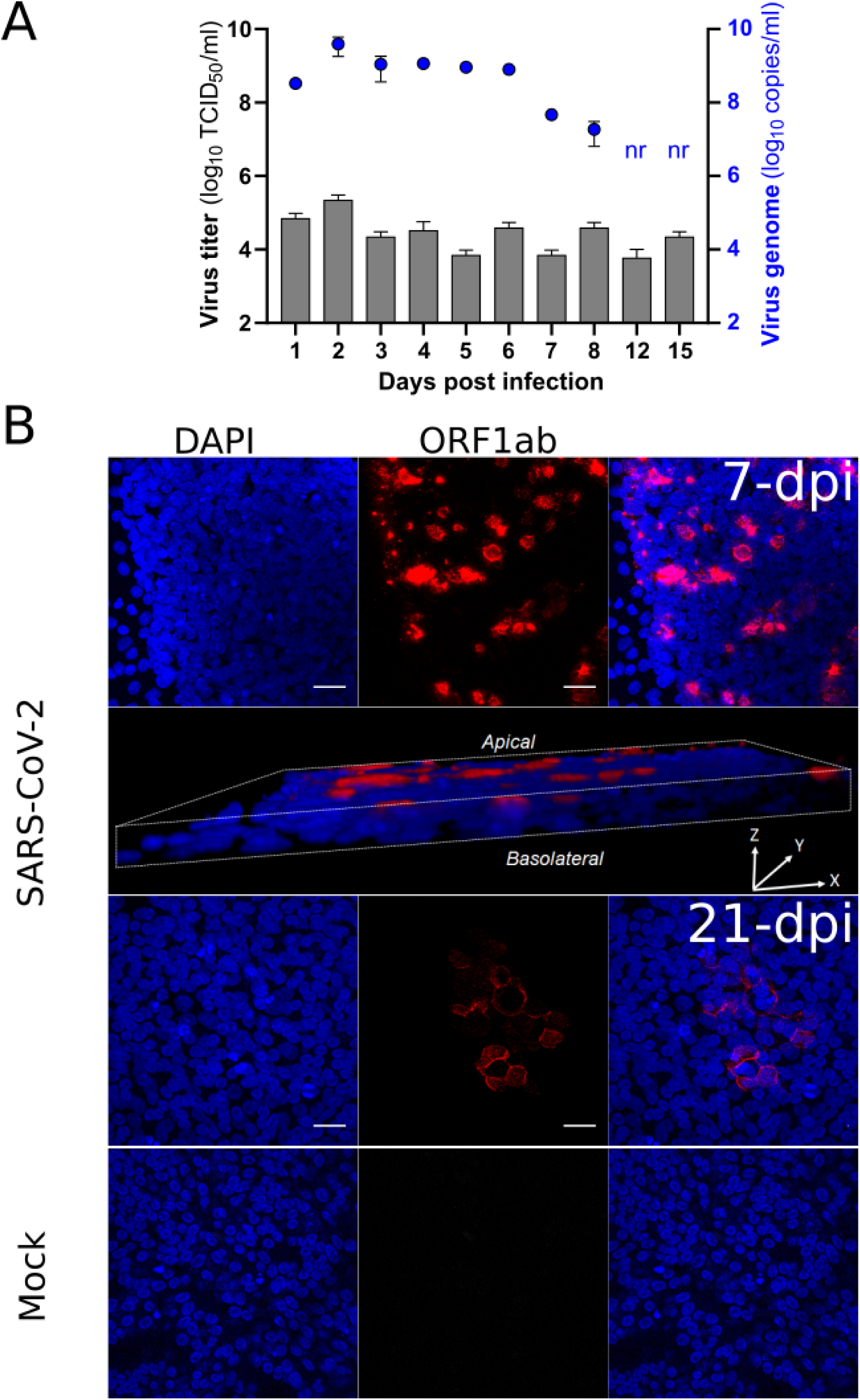
Persistent infection of nasal HAEEC by SARS-CoV-2. A) SARS-CoV-2 Omicron collected at the apical side of a duplicate (TCID_50_ means±SD). Virus titers (TCID_50_) are shown by bars (left y-axis) and virus genome copy numbers from RT-qPCR measurements are represented by blue dots (right y-axis). The virus genome copy numbers at days 12 and 15pi were not tested (nr). B) Intracellular presence of SARS-CoV-2 RNA (+) genome in HAEEC fixed at 7d and 21dpi, respectively. SARS-CoV-2 genomes (+) were stained by RNA fluorescence in situ hybridization (FISH) using oligonucleotide probes targeting the viral ORF1ab. The scale bar represents 30µm. A 3D projection computed by Fiji software is shown at 7dpi. Mock infected insert fixed at 21dpi and stained as the infected cells served as a negative control.

Together, these results support the notion that HAEEC can be infected with SARS-CoV-2 for periods of at least several weeks. This gives rise to a situation that resembles a persistent infection, where virus is steadily released without overt tissue death. Persistence in vivo may enhance the chance for viral recombination, which requires that two or more different infectious agents are present in the same cell. A survey of SARS-CoV-2 genomes from UK firstly observed mosaic genomes structures of Alpha VoC with other co circulating variants, likely to be from recombinant origin (**37**). Then, other potential SARS-CoV-2 recombinant lineage sequences were deposited on genomic databases and currently labelled XA to XL by the PANGO lineage nomenclature (**38**).

Similarly, as both the Delta and the Omicron VoC have been massively co-circulating in many countries across the world, it can be expected that Delta and Omicron recombinants will appear. In fact, this has recently been suggested in several independent preprints (**39**-**41**), acknowledged by the WHO with the BA.1 x AY.4 recombinant, nicknamed ‘Deltacron’, and labelled XD in the PANGO lineage nomenclature (**38**). Although it is presently unknown, if these recombinants will outcompete the known SARS-CoV-2 variants, virus recombinants raise concerns about increased transmission and pathogenicity. This issue requires continued monitoring, drug and vaccine development and explorations of repurposable drugs against COVID-19.

## Material and Methods

### Viruses

SARS-CoV-2 Omicron (B.1.1.529.1,BA.1; NH-RIVM-71076/2021) variant was obtained from the RIVM (Netherlands) through European Virus Archive global and expanded on VeroE6 cells grown in DMEM (Sigma Aldrich, Cat #D6429) supplemented with 4% of FCS (Gibco, Cat #10270-106), and 1X of NEAA (Sigma Aldrich, Cat #M7145). Virus titers were determined by TCID_50_ titration on VeroE6 cells according to the Spearman-Kärber method.

### Niclosamide

Niclo was purchased to Sigma Aldrich (cat#N3510-50G; lot #BCBD3349V) and solubilized according to manufacturer’s instructions.

### Nasal and bronchial HAEEC

Human nasal HAEEC (MucilAir™, Epithelix SA,Geneva, Switzerland) cultured on transwell inserts (24-well plate) were maintained at air-liquid interface according to the supplier’s instructionsand cell culture medium (Epithelix SA,Geneva, Switzerland, cat#EP05MM,). Nasal HAEEC from a pool of fourteen healthy donors (Batch nr: MP010).

Human bronchial were obtained from an individual donor (Donor 793, Epithelix SA, Geneva, Switzerland). Cells have been then seeded on Type IV collagen (Sigma Aldrich,cat#C5533) coated inserts (24-well plate) and amplified with PneumaCult™-Ex Basal Medium (Stemcell cat#05009) supplemented 1X with PneumaCult™-Ex 50X-Supplement (Stemcell, cat#05019) and hydrocortisone (Stemcell, cat#07925). Then cell have been differentiated with PneumaCult™-ALI Base Medium (Stemcell, cat#05002) supplemented with 1X of PneumaCult™-ALI 10X supplement (Stemcell, cat#05003), hydrocortisone (Stemcell, cat#07925), heparin (Stemcell, cat#07980), and 150 ng/mL of retinoic acid (Sigma Aldrich, cat#R2625).

### RNA FISH with branched DNA signal amplification

Inserts were fixed with 4% PFA in PBS for 30 min at RT, washed twice with PBS, dehydrated and permeabilized with absolute methanol overnight at -20°C. Samples were rehydrated by incubation with 75%, 50%, 25%, 0% of ice-cold methanol and PBST (Tween-20, 0.1%) for 5 min each. Rehydrated samples were washed on ice with a vol/vol solution of 5XSSCT /PBST for 5min, afterwards 5XSSCT alone for 5min. Samples were then FISH-stained against SARS-CoV-2 ORF1a mRNA using ViewRNA mRNA FISH assay according to the manufacturer’s instructions (ThermoFisher) with some modifications. Samples were hybridized with the SARS-CoV-2 targeting probes overnight at 40°C, then the subsequent pre-amplifier, amplifier, and Alexa Fluor 546 labeled probes hybridizations steps were done for 2h each at 40°C. SARS-CoV-2 targeting probes were custom-made #6007037-01 directed against the ORF1a sequences between positions 401-1327 (ThermoFisher). Subsequently, cells were incubated in PBS containing DAPI for 10 min at RT. Finally, the stained inserts have been detached from their plastic support, then slide/coverslip mounted, and then imaged using a SP8 confocal microscope (Leica).

### Phospolipidosis assay

Phospolipidosis assay was performed as described by Tummino and colleagues (**19**). Briefly, VeroE6 cells were seeded on a 96-well black imaging plate at a density of 15,000 cells per well and grown overnight in DMEM supplemented with 10%FCS and 1X NEAA. Then the medium was replaced with DMEM supplemented with 10%FCS, 1X NEAA, 7.5µM NBD-PE (ThermoFisher, cat#N360), and three concentrations of MB, MPA, Niclo, NH4Cl, and Amiodarone and incubated for 24hours at 37°C. Water and DMSO were used as solvent controls. Finally, cells were stained for 20 min at 37°C with a solution of DMEM supplemented with 100mM sodium pyruvate, 200mM L-Glutamine, 10% FCS, 1X NEAA, 10µg/mL Hoechst, and 2µM Ethidium homodimer-2 (EthD-2), and imaged with ImageXpress Micro confocal (Molecular Devices) microscope.

### SARS-CoV-2 infection of nasal and bronchial HAEEC tissue and drug treatment

This was carried out as published in Murer et al (**2**).

### RNA extraction and RT-qPCR

This was carried out as published in Murer et al (**2**).

## Acknowledgements

We acknowledge financial support from the Swiss National Science foundation (SNSF) to UFG (31CA30_196177/1) and a special Coronavirus Research grant from the University of Zurich to UFG. We are grateful to Dr. Adriano Aguzzi and Dr. Simone Hornemann for granting generous access to their BSL3 laboratory, and Dr. Maarit Suomalainen for discussions.

## Supplementary material

**Suppl Fig 1.**
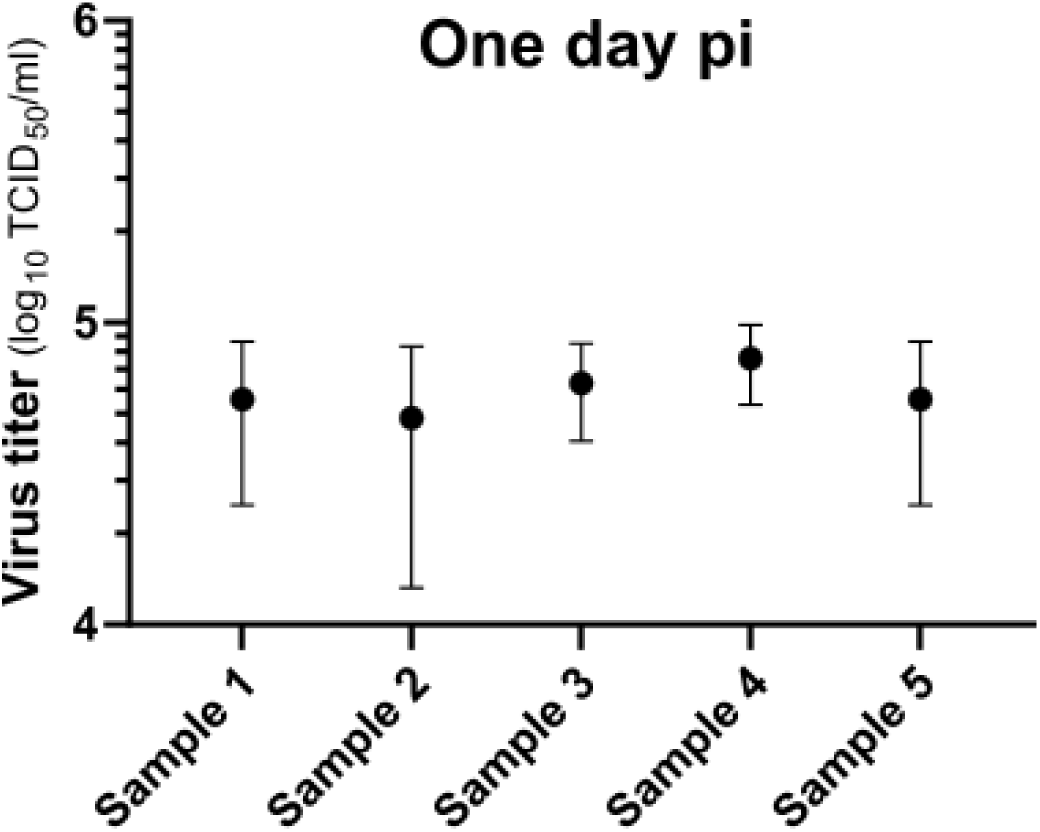
Apical infectious titer of SARS-CoV-2 Omicron in nasal HAEEC at one day pi. Data represent the means ± SD of three independent replicates of nasal HAEEC inoculated apically with 1,000 TCID_50_ units of SARS-CoV-2 Omicron.

**Suppl Fig 2.**
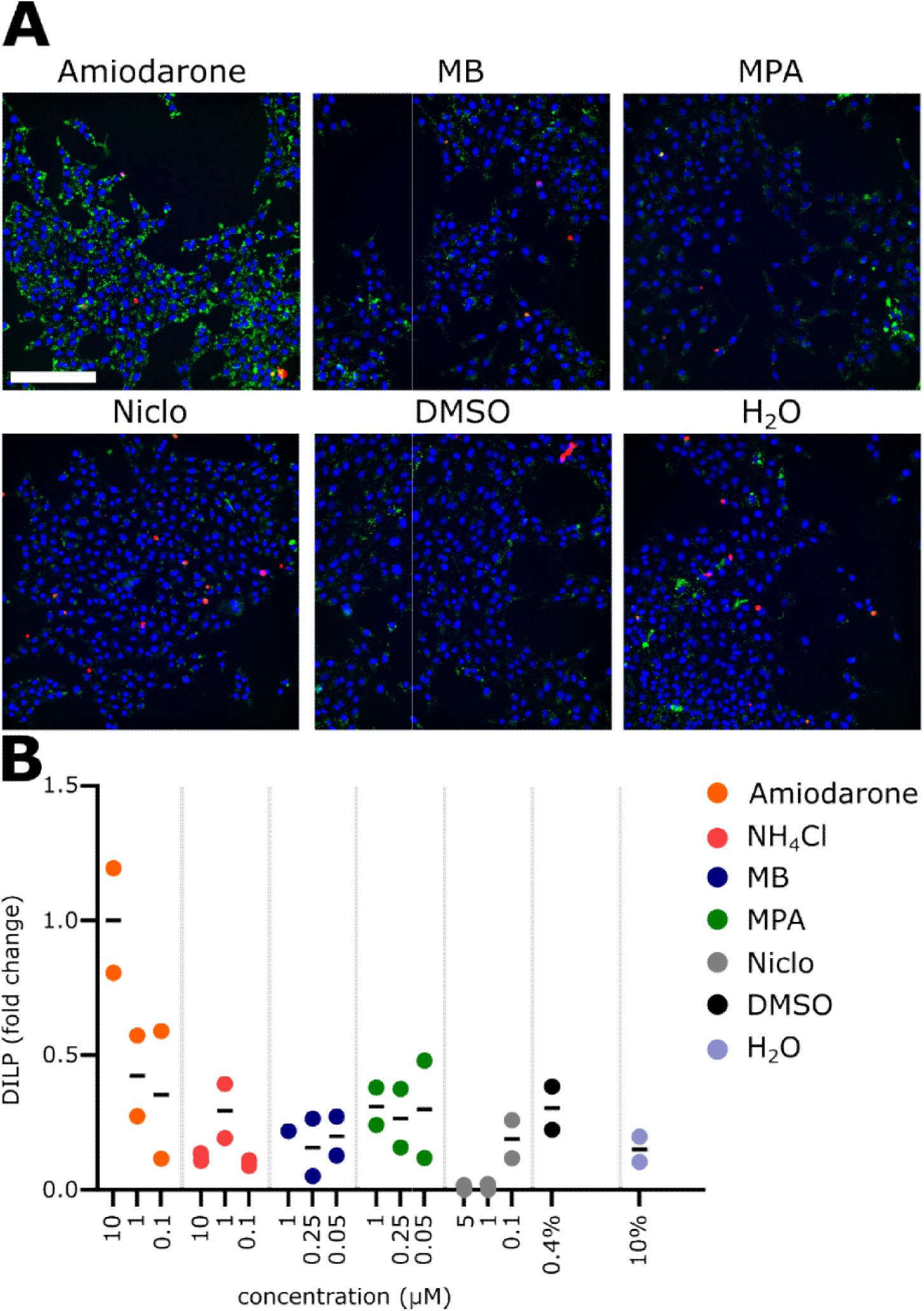
Drug-induced phospholipidosis. A) Example images of VeroE6 cells treated with the indicated drugs for 24 hours. Scale bar = 200 µm. B) Quantification of drug-induced phospholipidosis (DIPL). Fold change is calculated relative to the mean of the positive control Amiodarone.

## References

1. Weiss, A., Touret, F., Baronti, C., Gilles, M., Hoen, B., Nougairède, A., et al., 2021. Niclosamide shows strong antiviral activity in a human airway model of SARS-CoV-2 infection and a conserved potency against the Alpha (B.1.1.7), Beta (B.1.351) and Delta variant (B.1.617.2). PloS one. 16 (12), e0260958. https://doi.org/10.1371/journal.pone.0260958

2. Murer, L., Volle, R., Andriasyan, V., Petkidis, A., Gomez-Gonzalez, A., Yang, L., et al., 2022. Identification of broad anti-coronavirus chemical agents for repurposing against SARS-CoV-2 and variants of concern. Current Research in Virological Science. 3, 100019. https://doi.org/10.1016/j.crviro.2022.100019

3. World Health Organization, weekly epidemiological update on COVID-19 – 22 March 2022; https://www.who.int/publications/m

4. European Centre for Disease Prevention and Control (ECDC) SARS-CoV-2 variants dashboard, access 23 March 2022; https://www.ecdc.europa.eu/en/covid-19/situation-updates/variants-dashboard

5. Nextstrain, Genomic epidemiology of SARS-CoV-2 with global subsampling, access 23 March 2022; https://nextstrain.org/ncov/gisaid/global

6. Wei, C., Shan, K. J., Wang, W., Zhang, S., Huan, Q., Qian, W., 2021. Evidence for a mouse origin of the SARS-CoV-2 Omicron variant. Journal of genetics and genomics. 48 (12), 1111–1121. https://doi.org/10.1016/j.jgg.2021.12.003

7. Centers for Disease Control And Prevention. Science Brief: Omicron (B.1.1.529) Variant. 2021, access 23 March 2022; https://www.cdc.gov/coronavirus/2019-ncov/science/science-briefs/scientific-brief-omicron-variant.html

8. Iketani, S., Liu, L., Guo, Y., Liu, L., Chan, J. F., Huang, Y., et al., 2022. Antibody evasion properties of SARS-CoV-2 Omicron sublineages. Nature. 10.1038/s41586-022-04594-4. Advance online publication. https://doi.org/10.1038/s41586-022-04594-4

9. Allen, H., Tessier, E., Turner, C., Anderson, C., Blomquist, P., Simons, D., et al.,2022. Comparative transmission of SARS-CoV-2 Omicron (B.1.1.529) and Delta (B.1.617.2) variants and the impact of vaccination: national cohort study, England. medRxiv. https://doi.org/10.1101/2022.02.15.22271001

10. Rössler, A., Riepler, L., Bante, D., von Laer, D., Kimpel, J., 2022. SARS-CoV-2 Omicron Variant Neutralization in Serum from Vaccinated and Convalescent Persons. The New England journal of medicine. 386 (7), 698–700. https://doi.org/10.1056/NEJMc2119236

11. Lauring, A. S., Tenforde, M. W., Chappell, J. D., Gaglani, M., Ginde, A. A., McNeal, T., et al., 2022. Clinical severity of, and effectiveness of mRNA vaccines against, covid-19 from omicron, delta, and alpha SARS-CoV-2 variants in the United States: prospective observational study. BMJ (Clinical research ed.). 376, e069761. https://doi.org/10.1136/bmj-2021-069761

12. Netzl, A., Tureli, S., LeGresley, E., Mühlemann, B., Wilks, S. H., Smith, D. J., 2022. Analysis of SARS-CoV-2 Omicron Neutralization Data up to 2021-12-22. bioRxiv. https://doi.org/10.1101/2021.12.31.474032

13. Nyberg, T., Ferguson, N. M., Nash, S. G., Webster, H. H., Flaxman, S., Andrews, N., et al., 2022. Comparative analysis of the risks of hospitalisation and death associated with SARS-CoV-2 omicron (B.1.1.529) and delta (B.1.617.2) variants in England: a cohort study. Lancet (London, England). S0140-6736(22)00462-7. Advance online publication. https://doi.org/10.1016/S0140-6736(22)00462-7

14. Meng, B., Abdullahi, A., Ferreira, I., Goonawardane, N., Saito, A., Kimura, I., et al., 2022. Altered TMPRSS2 usage by SARS-CoV-2 Omicron impacts infectivity and fusogenicity. Nature. 603 (7902), 706–714. https://doi.org/10.1038/s41586-022-04474-x

15. Zhao, H., Lu, L., Peng, Z., Chen, L. L., Meng, X., Zhang, C., et al., 2022. SARS-CoV-2 Omicron variant shows less efficient replication and fusion activity when compared with Delta variant in TMPRSS2-expressed cells. Emerging microbes & infections. 11 (1), 277–283. https://doi.org/10.1080/22221751.2021.2023329

16. Hui, K., Ho, J., Cheung, M. C., Ng, K. C., Ching, R., Lai, K. L., et al., 2022. SARS-CoV-2 Omicron variant replication in human bronchus and lung ex vivo. Nature. 603 (7902), 715–720. https://doi.org/10.1038/s41586-022-04479-6

17. Kumar, S., Karuppanan, K., Subramaniam, G., 2022. Omicron (BA.1) and Sub-Variants (BA.1, BA.2 and BA.3) of SARS-CoV-2 Spike Infectivity and Pathogenicity: A Comparative Sequence and Structural-based Computational Assessment. bioRxiv. https://doi.org/10.1101/2022.02.11.480029

18. Shayman, J. A., & Abe, A., 2013. Drug induced phospholipidosis: an acquired lysosomal storage disorder. Biochimica et biophysica acta. 1831 (3), 602–611. https://doi.org/10.1016/j.bbalip.2012.08.013

19. Tummino, T. A., Rezelj, V. V., Fischer, B., Fischer, A., O’Meara, M. J., Monel, B., et al., 2021. Drug-induced phospholipidosis confounds drug repurposing for SARS-CoV-2. Science. 373 (6554), 541–547. https://doi.org/10.1126/science.abi4708

20. Andrews, P., Thyssen, J., Lorke, D., 1982. The biology and toxicology of molluscicides, Bayluscide. Pharmacology& therapeutics. 19 (2), 245–295. https://doi.org/10.1016/0163-7258(82)90064-x

21. Jurgeit, A., McDowell, R., Moese, S., Meldrum, E., Schwendener, R., Greber, U. F., 2012. Niclosamide is a proton carrier and targets acidic endosomes with broad antiviral effects. PLoS pathogens. 8 (10), e1002976. https://doi.org/10.1371/journal.ppat.1002976

22. Fonseca, B. D., Diering, G. H., Bidinosti, M. A., Dalal, K., Alain, T., Balgi, A. D., et al., 2012. Structure-activity analysis of niclosamide reveals potential role for cytoplasmic pH in control of mammalian target of rapamycin complex 1 (mTORC1) signaling. The Journal of biological chemistry. 287 (21), 17530–17545. https://doi.org/10.1074/jbc.M112.359638

23. Xu, J., Shi, P. Y., Li, H., Zhou, J., 2020. Broad Spectrum Antiviral Agent Niclosamide and Its Therapeutic Potential. ACS infectious diseases. 6 (5), 909– 915. https://doi.org/10.1021/acsinfecdis.0c00052

24. Kaidashev, I., Shlykova, O., Izmailova, O., Torubara, O., Yushchenko, Y., Tyshkovska, T., et al., 2021. Host gene variability and SARS-CoV-2 infection: A review article. Heliyon. 7 (8), e07863. https://doi.org/10.1016/j.heliyon.2021.e07863

25. Suomalainen, M., Greber, U. F., 2021. Virus Infection Variability by Single-Cell Profiling. Viruses. 13 (8), 1568. https://doi.org/10.3390/v13081568

26. Pereira, N. L., Ahmad, F., Byku, M., Cummins, N. W., Morris, A. A., Owens, A., et al., 2021. COVID-19: Understanding Inter-Individual Variability and Implications for Precision Medicine. Mayo Clinic proceedings. 96 (2), 446–463. https://doi.org/10.1016/j.mayocp.2020.11.024

27. Hang, J., Chen, H., Tian, P., Yu, R., Wang, M., Zhao, M., 2022. Preparation and pharmacodynamics of niclosamide micelles.Journal of Drug Delivery Science and Technology. 68, 103088. https://doi.org/10.1016/j.jddst.2021.103088.

28. Cairns, D. M., Dulko, D., Griffiths, J. K., Golan, Y., Cohen, T., Trinquart, L., et al., 2022. Efficacy of Niclosamide vs Placebo in SARS-CoV-2 Respiratory Viral Clearance, Viral Shedding, and Duration of Symptoms Among Patients With Mild to Moderate COVID-19: A Phase 2 Randomized Clinical Trial. JAMA network open. 5 (2), e2144942. https://doi.org/10.1001/jamanetworkopen.2021.44942

29. Backer, V., Sjöbring, U., Sonne, J., Weiss, A., Hostrup, M., Johansen, H. K., et al., 2021. A randomized, double-blind, placebo-controlled phase 1 trial of inhaled and intranasal niclosamide: A broad spectrum antiviral candidate for treatment of COVID-19. The Lancet regional health. Europe, 4, 100084. https://doi.org/10.1016/j.lanepe.2021.100084

30. Rambaut, A., Loman, N., Pybus, O. G., Barclay, W., Barret, J., Carabelli, A., et al., 2020. Preliminary genomic characterisation of an emergent SARS-CoV-2 lineage in the UK defined by a novel set of spike mutations. https://virological.org/t/preliminary-genomic-characterisation-of-an-emergent-sars-cov-2-lineage-in-the-uk-defined-by-a-novel-set-of-spike-mutations/563

31. Chaloner Larsson, G., Johnson-Lussenburg, C. M., 1981. Establishment and maintenance of a persistent infection of L132 cells by human coronavirus strain 229E. Archives of virology. 69 (2), 117–129. https://doi.org/10.1007/BF01315155

32. Arbour, N., Côté, G., Lachance, C., Tardieu, M., Cashman, N. R., Talbot, P. J. 1999. Acute and persistent infection of human neural cell lines by human coronavirus OC43. Journal of virology. 73 (4), 3338–3350. https://doi.org/10.1128/JVI.73.4.3338-3350.1999

33. Maponga, T. G., Jeffries, M., Tegally, H., Sutherland, A. D., Wilkinson, E., Lessels, R., et al., 2022. Persistent SARS-CoV-2 infection with accumulation of mutations in a patient with poorly controlled HIV infection. Available at SSRN: https://ssrn.com/abstract=4014499 or http://dx.doi.org/10.2139/ssrn.4014499

34. Choi, B., Choudhary, M. C., Regan, J., Sparks, J. A., Padera, R. F., Qiu, X., et al., 2020. Persistence and Evolution of SARS-CoV-2 in an Immunocompromised Host. The New England journal of medicine. 383 (23), 2291–2293. https://doi.org/10.1056/NEJMc2031364

35. Bianco, A., Capozzi, L., Del Sambro, L., Simone, D., Pace, L., Rondinone, V., et al., 2022. Persistent SARS-CoV-2 Infection in a Patient With Non-hodgkin Lymphoma: Intra-Host Genomic Diversity Analysis. Frontiers in Virology. 2, 758191. https://doi.org/10.3389/fviro.2022.758191

36. Vibholm, L. K., Nielsen, S., Pahus, M. H., Frattari, G. S., Olesen, R., Andersen, R., et al., 2021. SARS-CoV-2 persistence is associated with antigen-specific CD8 T-cell responses. EBioMedicine 64, 103230. https://doi.org/10.1016/j.ebiom.2021.103230

37. Jackson, B., Rambaut, A., Pybus, O. G., Robertson, D.L., Connor, T., Loman, N. J., et al., 2021. Recombinant SARS-CoV-2 genomes involving lineage B.1.1.7 in the UK. https://virological.org/t/recombinant-sars-cov-2-genomes-involving-lineage-b-1-1-7-in-the-uk/658

38. Rambaut, A., Holmes, E. C., O’Toole, Á., Hill, V., McCrone, J. T., Ruis, C., et al., 2020. Nature Microbiology. 5, 1403–1407.https://doi.org/10.1038/s41564-020-0770-5

39. Lacek, K. A., Rambo-Martin, B. L., Batra, D., Zheng, X., Sakaguchi, H., Peacock, T., et al., 2022. Identification of a Novel SARS-CoV-2 Delta-Omicron Recombinant Virus in the United States. bioRxiv. https://doi.org/10.1101/2022.03.19.484981

40. Bolze, A., White, S., Basler, T., Dei Rossi, A., Roychoudhury, P., Alexander L. et al., 2022. Evidence for SARS-CoV-2 Delta and Omicron co-infections and recombination. medRxiv. https://doi.org/10.1101/2022.03.09.22272113

41. Colson, P., Fournier, P. E., Delerce, J., Million, M., Bedotto, M., Houhamdi, L., et al., 2022. Culture and identification of a “Deltamicron” SARS-CoV-2 in a three cases cluster insouthern France. medRxiv. https://doi.org/10.1101/2022.03.03.22271812

